# Whole genome-based Characterization of Virulence and Antimicrobial Resistance Determinants in Clinical *Campylobacter jejuni* Isolates from Minnesota, 2018-2021

**DOI:** 10.64898/2026.04.14.718608

**Authors:** Haejin Hwang, Melanie Orth, Selina Jawahir, Annastasia Gross, Xiong Wang, Dave Boxrud, Kirk Smith, Jisun Haan

**Author notes:** Corresponding author: Jisun Haan.

## Abstract

*Campylobacter jejuni* is a leading cause of foodborne gastroenteritis globally and is classified by the CDC as a serious public health threat due to increasing resistance to fluoroquinolones and macrolides. This study used whole-genome sequencing to characterize the virulence and antimicrobial resistance profiles of 2,783 clinical *C. jejuni* isolates collected from Minnesota residents from 2018 through 2021. More than 90% of the isolates had genes related to stress defense (*rpoN* and *htrB*), cytolethal distending toxin (*cdtA, cdtB*, and *cdtC*), and cell adhesion and invasion (*ciaB, cadF*, and *flaC*). A diverse array of antimicrobial resistance genes was detected, with beta-lactam resistance genes having a particularly high prevalence. The *gyrA* point mutation associated with quinolone resistance was present in 29% of isolates. To evaluate the correlation between genotypic and phenotypic antimicrobial resistance profiles, the antimicrobial susceptibility testing results from a subset of isolates were compared with genotypic resistance profiles. Results showed a strong overall correlation, particularly for tetracycline and quinolones, though 24 discrepancies were detected. In the majority of discrepancies (n=21), genomic antimicrobial resistance markers were absent in isolates that were phenotypically resistant, suggesting possible unknown resistance mechanisms or limitations in current sequencing methods. The remaining three discrepancies occurred in isolates that had the *tet*(O) resistance gene but were susceptible to tetracycline phenotypically. These findings highlight the value of whole genome sequencing in improving antimicrobial resistance surveillance and understanding virulence factors in *C. jejuni*, supporting its integration into routine monitoring practices to better manage and understand antimicrobial resistance in foodborne pathogens.

## Introduction

*Campylobacter jejuni* is one of the leading causes of bacterial gastroenteritis in humans globally (1). In healthy individuals, campylobacteriosis is typically self-limiting (2). However, severe infections can occur, especially in at-risk populations, such as children, the elderly, and the immunocompromised (3). In such cases, antibiotic treatment with macrolides or fluoroquinolones is often necessary (3). There is a growing concern over the increasing resistance of *C. jejuni* to antibiotics worldwide (4, 5), which poses significant public health risks. The National Antimicrobial Resistance Monitoring System (NARMS) 2019 Integrated Report indicated that the percentage of fluroquinolone-resistant *C. jejuni* in human isolates rose from 29% in 2018 to 34% in 2019 (6). Similar increases in resistance were observed in chicken retail samples and cecal contents (6), reducing the effectiveness of fluroquinolone treatment for campylobacteriosis.

The natural reservoirs of *C. jejuni* include a wide range of wild and domesticated animals, including wild birds, cattle, sheep, and poultry. Consumption of undercooked or raw poultry products is considered the major source of *C. jejuni* in sporadic campylobacteriosis in humans (7), while consumption of unpasteurized milk or contaminated water is often linked to outbreaks (8).

Pulsed-field gel electrophoresis (PFGE) was considered a gold standard method for foodborne bacterial pathogen subtyping performed by public health laboratories (9). In 2019, the PulseNet National Laboratory Network transitioned from PFGE to whole-genome sequencing (WGS) for the detection and investigation of nationwide foodborne disease outbreaks in the United States, improving discrimination and precision of investigations (9). WGS can generate high-resolution genetic information with relatively quick turn-around time at a reasonable cost. The sequencing data generated by WGS allows comprehensive phylogenetic analysis of a set of bacterial strains, as well as screening of molecular markers of virulence and antimicrobial resistance.

The Minnesota Department of Health, Public Health Laboratory (MDH-PHL) has performed routine WGS of all available *Campylobacter* isolates since 2018 to aid in outbreak detection and investigation. However, extensive genomic characterization of these isolates has not been conducted. In this study, we used WGS to provide comprehensive genotypic antimicrobial resistance (AMR) and virulence factor profiles. Phenotypic resistance data from a subset of clinical isolates was compared to the genotypic AMR profiles to evaluate the accuracy of WGS in predicting phenotypic antimicrobial resistance in *C. jejuni*.

## Materials and Methods

### Bacterial isolate collection

*Campylobacter* is a reportable pathogen according to Minnesota Reportable Disease Rule, 5605.7040. As a part of Foodborne Diseases Active Surveillance Network (FoodNet) activities, MDH conducts active surveillance to ascertain 100% of laboratory-confirmed *Campylobacter* infections in Minnesota residents. Specimens that test *Campylobacter*-positive by culture independent diagnostic testing (CIDT) and *Campylobacter* isolates must be submitted to the MDH-PHL for all cases of campylobacteriosis; the MDH-PHL attempts to isolate *Campylobacter* from all submitted positive clinical specimens and identifies all *Campylobacter* isolates to species. The *C. jejuni* isolates included in this study comprised all 3,628 clinical *C. jejuni* isolates collected from Minnesota residents from 2018 through 2021.

### Isolation and identification of *Campylobacter jejuni*

When a clinical laboratory submitted a positive clinical specimen to the MDH-PHL, *Campylobacter* spp. isolation was attempted from stool samples or rectal swabs submitted in Cary-Blair or other enteric transport media using a filtration method at the MDH-PHL. A cellulose acetate filter (0.65µm) was placed on two *Campylobacter* blood agar (CBA) plates, and 6-8 drops of stool sample were directly placed on the filter. After incubating the plate for 45 minutes to 1hr at 36ºC, the filter was removed. One plate was incubated at 42ºC for 24-72 hrs under microaerophilic conditions (87.4% N_2_, 7.6% CO_2_, 5% O_2_) and the other was incubated at 36 ºC for 48 hours under increased hydrogen microaerophilic condition (81.3% N2, 8.6% CO2, 7.1% H2,3% O2). The clinical specimen was also directly plated to Cefoperazone, Vancomycin, Amphotericin B Agar (CVA) plate for isolation, and incubated at 42ºC for 24-72 hrs under microaerophilic conditions.

An oxidase test was performed on *Campylobacter*-like colonies. Colonies with a positive oxidase test were sub-cultured to CBA and incubated for 24 hrs at 42ºC under microaerophilic conditions. Gram’s stain and matrix-assisted laser desorption/ionization-time of flight Mass Spectrometry (MALDI-TOF MS) identification (Bruker, USA) were performed to determine the species of the *Campylobacter* isolate.

Gram’s stain, oxidase and MALDI-TOF MS identification were performed to confirm the species when *C. jejuni* isolates were submitted instead of clinical specimens.

### Whole genome sequencing and genome assembly

*Campylobacter* genomic DNA was extracted on QIAcube Classic using the QIAamp DNA Mini QIAcube kit (Qiagen, Germany) or MagNA Pure LC (Roche, Switzerland) using the Pure LC Total Nucleic Acid (TNA) Isolate kit (MagNA Pure) following the manufacturer’s instructions. DNA concentration was measured using the Quant-iT dsDNA HS Assay kit (Invitrogen, USA). Libraries were prepared using the Nextera XT index kit (Illumina, USA) following the manufacturer’s instructions. Sequencing was performed on the Illumina Miseq platform with 250 bp-paired-end reads using V2 chemistries. Raw reads were trimmed and quality-filtered using Trimmomatic PE (10). High-quality reads were *de novo* assembled by Skesa (ver. 2.4.0) using default parameters (11). Assembly metrics were calculated using QUAST (ver. 5.0.2) with *C. jejuni* NCTC 11168 as a reference genome (12). BUSCO (ver. 5.2.2) was performed to assess genome assembly completeness (13). Lastly, average genome coverage was calculated using Samtools (ver. 1.15) (14). Assembled genomes were excluded from further analysis using the following exclusion criteria: 1) N50 < 10,000; 2) total length of assembled genome > 2,200,000 bps; 3) genome completeness by BUSCO < 95%; and 4) average genome coverage < 20X. The final dataset used for the downstream analyses consisted of 2,783 genomes.

### WGS analysis: AMR prediction and virulence factor screening

The presence of acquired resistance genes and chromosomal point mutations were screened using ResFinder (ver. 4.0) and BLASTn (ver. 2.9.0) (15, 16). The minimum identity threshold for matching was set to 90% with 60% minimum coverage. Virulence factor screening was performed using ABRicate (ver. 1.0.1) against the Virulence Factor Database (VFDB) using the default parameters (17, 18). A virulence factor was considered present if the minimum proportion of the gene covered (coverage) was greater than 90%. Specifically, the distribution of 17 virulence genes associated with cytolethal distending toxin (CDT) producing genes (*cdtA, cdtB*, and *cdtC*), cell invasion and adhesion (*ciaB, ciaC, flaC, cadF*, and *jlpA*), motility (*flaA* and *flaB*), stress response (*rpoN* and *htrB*), and Type IV secretion system (*virB8, virB9, virB10, virB11*, and *virD4*) were investigated.

### Antimicrobial susceptibility testing

A subset (n=547) of *C. jejuni* isolates from the MDH-PHL was selected to measure minimum inhibitory concentration (MIC) using broth microdilution. These clinical isolates were selected randomly, with every 20^th^ isolate submitted for antimicrobial susceptibility testing (AST) as part of the National Antimicrobial Resistance Monitoring System (NARMS). Susceptibility to nine antibiotics was measured: azithromycin, ciprofloxacin, erythromycin, tetracycline, florfenicol, nalidixic acid, telithromycin, clindamycin, and gentamicin. Briefly, 3-5 isolated colonies were suspended in 5 mL of Mueller Hinton broth with TES (Thermo Fisher Scientific, USA). The suspension was adjusted to 0.5 McFarland turbidity, and 100 µL of the suspension was diluted in 11 mL Mueller Hinton broth with lysed horse blood (Thermo Fisher Scientific, USA). Sensititre™ *Campylobacter* CMVCAMPY panels (Thermo Fisher Scientific, USA) were inoculated with 100 µL of the inoculum using TREK Sensititre AIM™ Automated Inoculation Delivery System (TREK Diagnostic Systems, USA) with *C. jejuni* ATCC 33560 as a quality control strain. Panels were incubated at 37ºC for 48 hrs under microaerophilic conditions. Panels were manually read using Sensititre Vizion Digital MIC viewing system (TREK Diagnostic Systems, USA), and MICs were interpreted as susceptible or resistant using clinical breakpoints or epidemiological cutoff values (ECOFF) from the European Committee on Antimicrobial Susceptibility Testing (EUCAST) (19).

### Phenotypic and genotypic AMR profile comparison

The Cohen’s Kappa statistic was calculated to assess the level of concordance between the resistance phenotype obtained from the broth microdilution method and the genotype obtained using ResFinder in RStudio (ver. 4.1.2) (20). Resistance genotype was defined as the presence of one or more resistance genes and/or point mutations for each antimicrobial tested using the broth microdilution method.

## Results

### Genome characteristics

A total of 3,628 *C. jejuni* isolates from human clinical cases collected in Minnesota from 2018 through 2021 were initially obtained for the study. Following the application of filtering criteria, including genome completeness, average genome coverage, total genome length, and N50 of the assembled genome, 2,783 isolates were included in the final analysis (Table S1). Of these, 912 isolates were collected in 2018, 739 in 2019, 484 in 2020, and 648 in 2021. The majority of the isolates (n=2,739) were from stool or rectal swabs, with 10 from blood and 34 from other specimen types.

The average GC content of the compiled genomes was 30.4% (range: 29.9 – 31.4%). The average length of assembled genomes was 1,677,537 bps (range: 1,533,180 - 1,961,390 bps) with an average genome completeness of 98.8% (range: 95.1 – 99.8%). The average genome coverage was 57.8X (range: 20.1 – 558.01X).

### Virulence factor screening

Two virulence genes, *cadF* and *flaC*, which are involved in cell adhesion and invasion, were present in all 2,783 isolates (Table 1). The Cytolethal Distending Toxin (CDT)-producing genes (*cdtA, cdtB*, and *cdtC*), stress defense genes (*rpoN* and *htrB*), and two cell adhesion and invasion genes (*ciaC* and *jlpA*) were present in more than 95% of the isolates, while *ciaB* was found in 75% of the isolates. Approximately a quarter of isolates carried *flaA* and *flaB*, while less than 6% of the isolates carried the Type IV secretion system virulence genes.

**Table 1.**
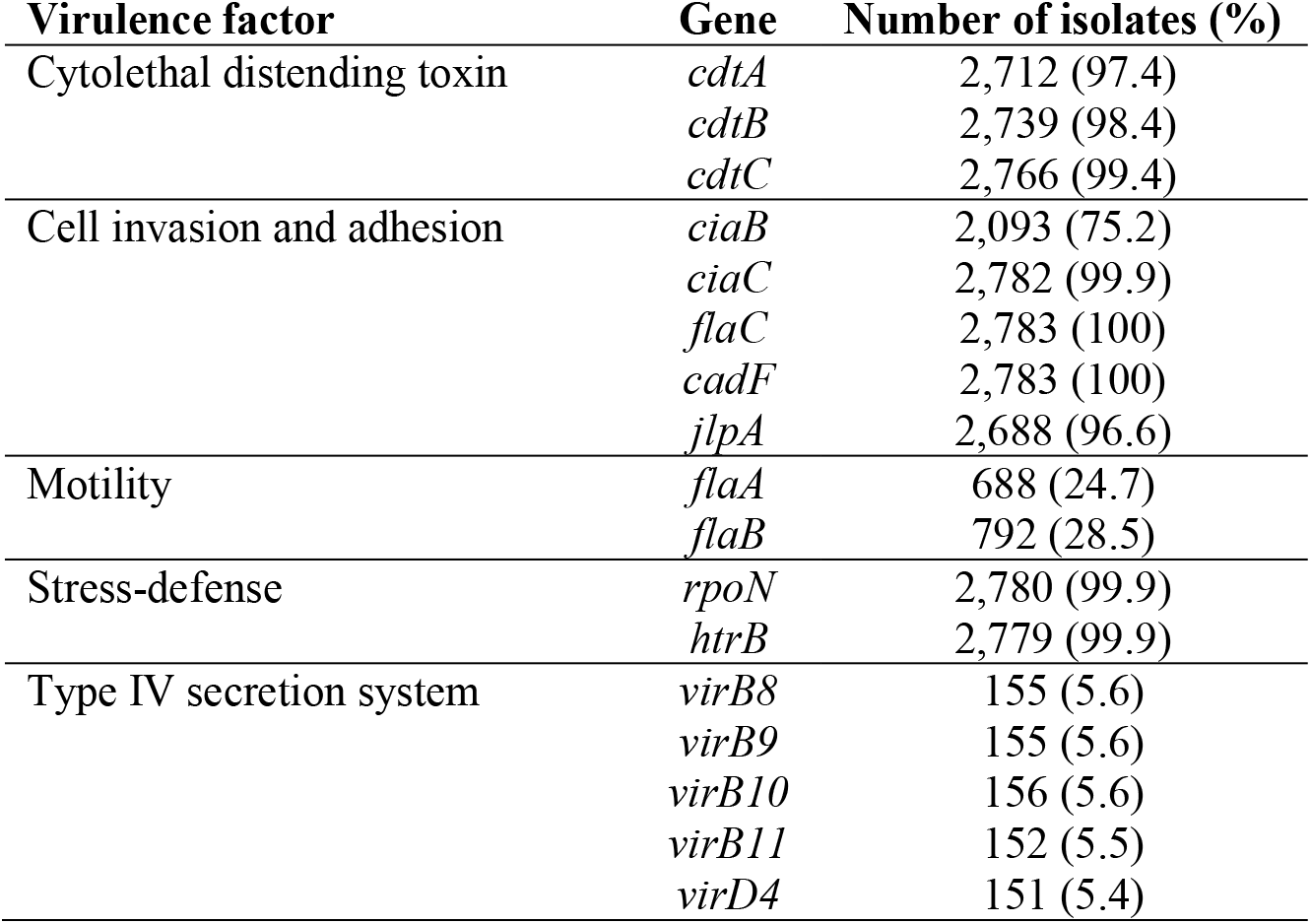
Prevalence of virulence genes in 2,783 clinical *C. jejuini* isolates from Minnesota residents, 2018– 2021.

### Genotypic antimicrobial resistance

ResFinder was run on 2,783 *C. jejuni* isolates to identify antimicrobial resistance genes and point mutations (Tables S2 and S3). Twenty-five acquired antimicrobial resistance genes and nine chromosomal point mutations were identified. Distribution of acquired antimicrobial resistance genes and chromosomal point mutations are presented in Table 2 and Table 3, respectively. The majority of the isolates (n=2571, 92.4%) had at least one acquired resistance gene or point mutation associated with antimicrobial resistance, while 212 (7.6%) were genotypically pan-susceptible. Among the isolates with antimicrobial resistance determinants, 2520 (90.5%) had at least 1 acquired resistance gene, and 837 (29.7%) had at least one point mutations associated with antimicrobial resistance.

**Table 2.** Distribution of antimicrobial resistance genes in clinical *C. jejuni* isolates (n=2,520) from Minnesota residents, 2018 – 2021.

| Drug Class | Antimicrobial resistance gene | Number of isolates with gene (%)<br>(n=2,520) |
| --- | --- | --- |
| Macrolides | *ermB* | 1 (0.04) |
| Lincosamides | *lnuC* | 1(0.04) |
| Tetracyclines | *tet*(O) | 1331 (52.8) |
|  | *tet*(O/32/O) | 81 (3.2) |
|  | *tet*(O/W/32/O) | 2 (0.08) |
| Beta-lactams | *bla*<sub>OXA61</sub> | 1,617 (64.2) |
|  | *bla*<sub>OXA184</sub> | 96 (3.8) |
|  | *bla*<sub>OXA185</sub> | 14 (0.6) |
|  | *bla*<sub>OXA193</sub> | 1,533(60.8) |
|  | *bla*<sub>OXA446</sub> | 2 (0.08) |
|  | *bla*<sub>OXA447</sub> | 31 (1.2) |
|  | *bla*<sub>OXA448</sub> | 129 (5.1) |
|  | *bla*<sub>OXA449</sub> | 45 (1.8) |
|  | *bla*<sub>OXA450</sub> | 1,472 (58.4) |
|  | *bla*<sub>OXA451</sub> | 1,470 (58.4) |
|  | *bla*<sub>OXA453</sub> | 1,481 (58.8) |
|  | *bla*<sub>OXA460</sub> | 129 (5.1) |
|  | *bla*<sub>OXA461</sub> | 145 (5.8) |
|  | *bla*<sub>OXA465</sub> | 41 (1.6) |
|  | *bla*<sub>OXA466</sub> | 34 (1.3) |
|  | *bla*<sub>OXA489</sub> | 1,474 (58.5) |
| *Aminoglycosides* | *aph*3'-III | 278 (11) |
|  | *aph*(2'')-Ih | 13 (0.5) |
|  | *aph*(2'')-If | 7 (0.3) |
|  | *ant*(6')-Ia | 26 (1) |

**Table 3.** Distribution of point mutations in clinical *C. jejuni* isolates from Minnesota residents, 2018 – 2021.

| Gene | Point mutation | Number of isolates with mutation (%) (n=837) |
| --- | --- | --- |
| *gyrA* (n=821) | T86I | 819 (97.8) |
|  | D90N | 6 (0.7) |
|  | P104S | 2 (0.2) |
|  | T86A | 1 (0.1) |
|  | T86V | 1 (0.1) |
| 23S rRNA (n=53) | S2075AG | 41 (5.0) |
|  | S2074AT | 12 (1.4) |
| *rpsL* (n=4) | K43R | 2 (0.2) |
|  | K88R | 2 (0.2) |

Among the 25 acquired resistance genes, *bla*_OXA-61_ (n=1617; 64.2%) was, by far, the most frequently identified. Multiple *bla*_OXA_ genes, which are involved in encoding β-lactamase, were identified in individual isolates. In addition to *bla*_OXA-61,_ the genes *bla*_OXA-489_, *bla*_OXA-453_, *bla*_OXA-451_, *bla*_OXA-450_, and *bla*_OXA-193_ were also commonly detected (Table 2). The second most frequently observed resistance gene was *tet*(O) (n=1331, 52.8%). In addition to *tet*(O), the mosaic tetracycline genes *tet*(O/32/O) (n=81, 3.2%) and *tet*(O/W/32/O) (n=2, 0.08%) were also identified, though at a lower frequency. Four resistance genes associated with aminoglycosides were identified with *aph(3-’)-III* (n=278; 11%) being the most frequently observed aminoglycoside resistance genes among the isolates. Notably, *aph(2’’)-If*, which has been reported to confer resistance to kanamycin and gentamicin in *C. jejuni* (21), was detected in seven isolates.

The most common point mutations were *gyrA* point mutations associated with fluoroquinolone resistance (n=821, 98.1%) (Table 3). The T86I (n=819, 97.8%) mutation was the most common *gyrA* point mutation, followed by D90N (n=6, 0.7%), P104S (n=2, 0.2%), T86A (n=1, 0.1%) and T86V (n=1, 0.1%). Notably, the less common *gyrA* point mutation, D90N and P104S, were detected together with T86I (n=6, n=2, respectively). A total of 53 (6.3%) *C. jejuni* isolates had point mutations in the 23S rRNA that confer resistance to azithromycin, erythromycin, clindamycin, and telithromycin. Four isolates had a point mutation in the *rpsL* gene which confers resistance to streptomycin.

### Phenotypic antimicrobial resistance

MICs of a subset of clinical *C. jejuni* isolates (n=547) is presented in Figure 1. These isolates were randomly selected to test MICs as a part of NARMS surveillance. Two hundred eighteen (40%) of these were pan-susceptible to all antibiotics tested. The most frequently observed resistance phenotype was tetracycline (n=286, 52.3%), followed by quinolones (n=168, 30.7%), macrolides (n=13, 2.4%), lincosamides (n=13, 2.4%), aminoglycosides (n=10, 1.8%), and phenicols (n=4, 0.7%). One isolate was resistant to ciprofloxacin but susceptible to nalidixic acid.

**Figure 1.**
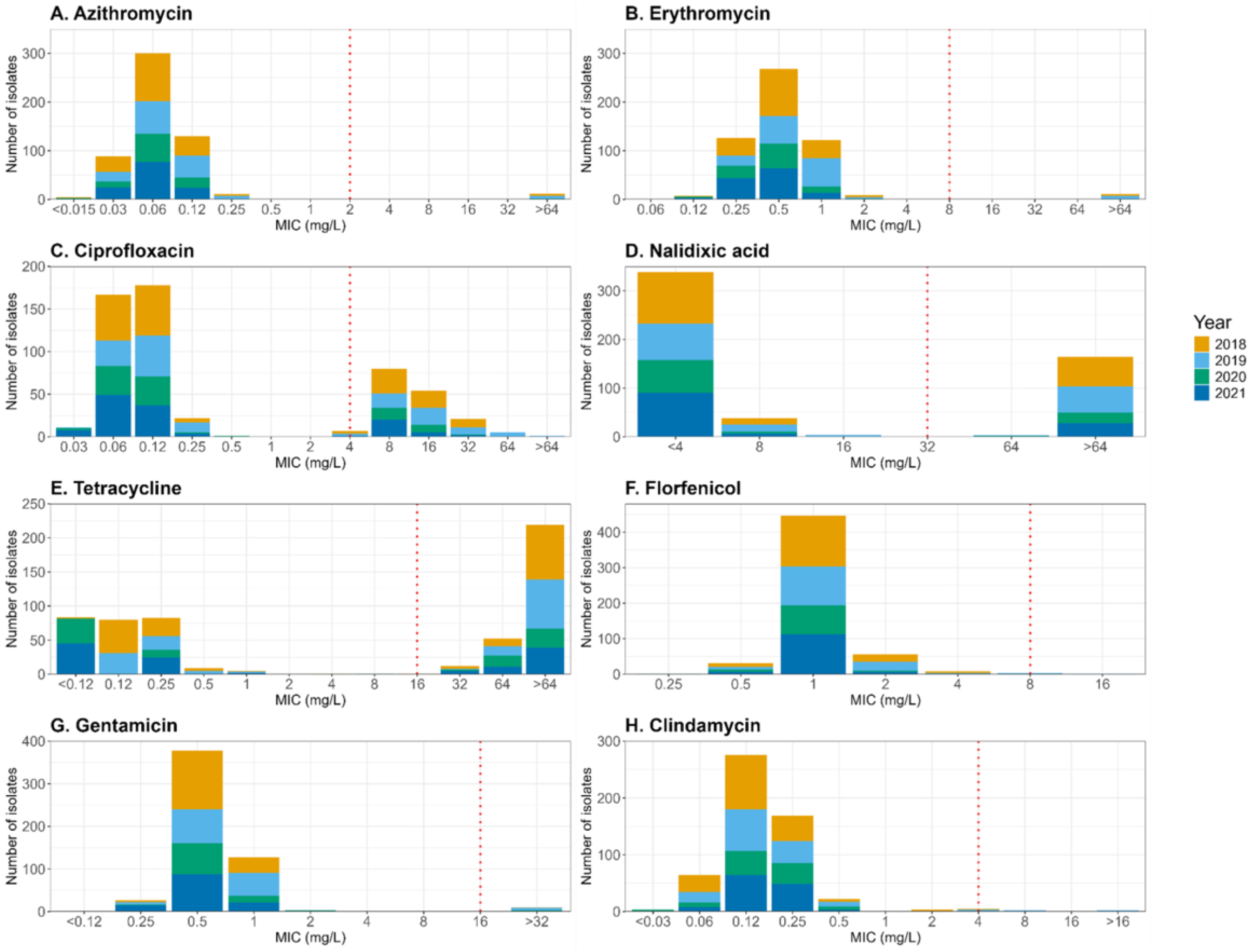
Distribution of MIC values for 8 antibiotics from a subset of clinical *C. jejuni* isolates from Minnesota residents, 2018-2021. *The breakpoint concentration is denoted by a red dotted line.

Nineteen isolates (3.5%) were multidrug resistant (MDR) (resistant to 3 or more antibiotic classes) (Figure 2). The most common MDR pattern was AZM-CIP-CLI-ERY-NAL-TET (n=6). Five MDR isolates were resistant to five classes of antibiotics (macrolides, quinolones, aminoglycosides, lincosamides, and tetracyclines).

**Figure 2.**
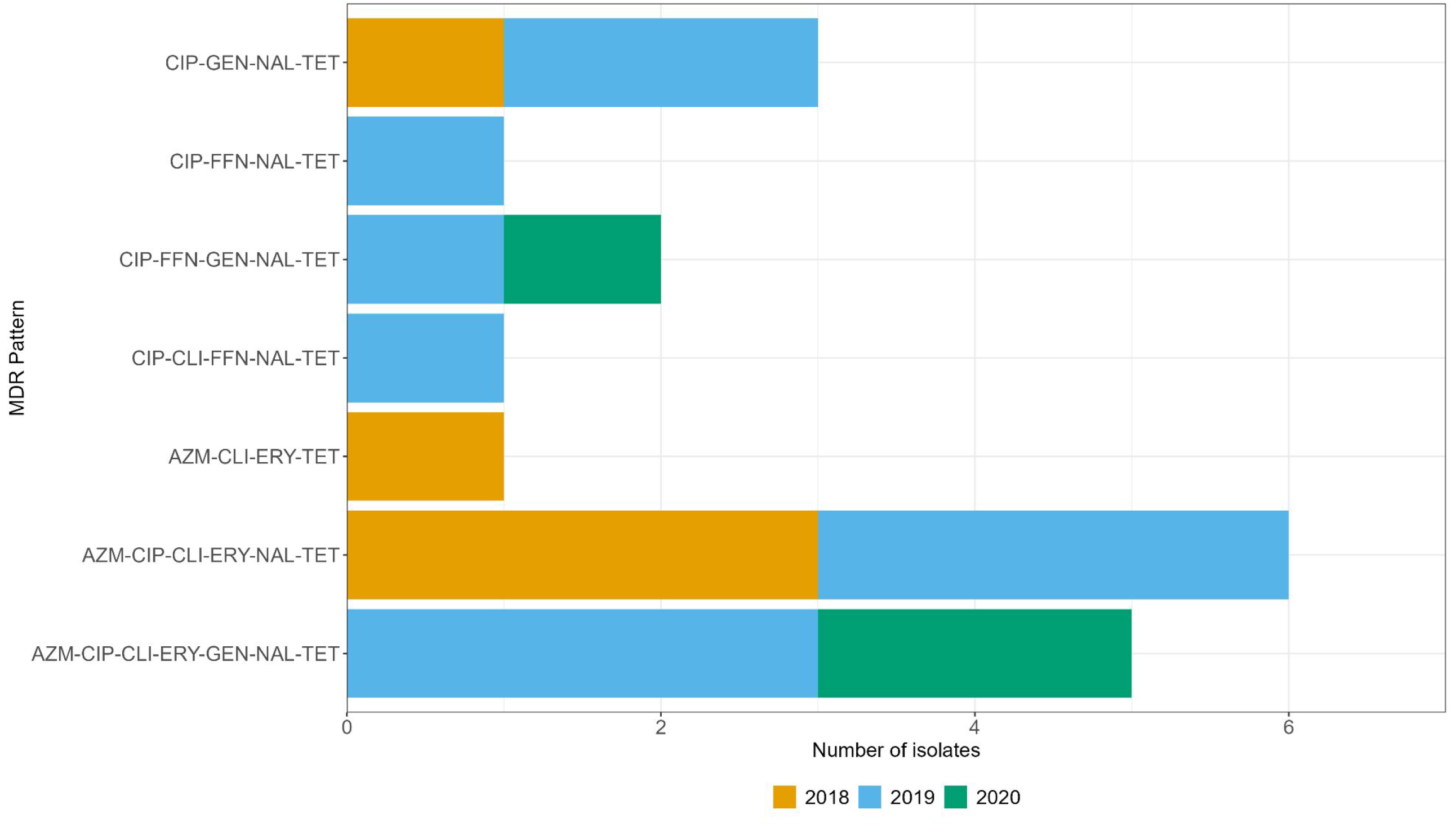
Distribution of multidrug resistance patterns among a subset of clinical *C. jejuni* isolates from Minnesota residents, 2018-2021. Abbreviations: CIP, ciprofloxacin; GEN, gentamicin; NAL, nalidixic acid; TET, tetracycline; FFN, florfenicol; CLI, clindamycin; AZM, azithromycin; ERY, erythromycin.

### Comparison between phenotypic and genotypic antimicrobial resistance profiles

Overall, high concordance between phenotypic resistance and genotypic resistance was observed across all antibiotic classes tested, with the average Cohen’s Kappa statistic of 0.92 (Table 4). A total of 24 mismatches were observed, and mismatches occurred across all antibiotic classes. A majority (n=21) of the mismatches occurred because isolates were phenotypically resistant to the antibiotics but lacked resistance genes or point mutations that conferred the phenotypic resistance. This type of mismatch was most frequently observed with tetracycline (n=11), followed by quinolones (n=5), lincosamides (n=3), aminoglycosides (n=3), and macrolides (n=2). The other three mismatches occurred in the isolates that carried *tet*(O) but were phenotypically susceptible to tetracycline with their MICs against tetracycline below 1 µg/mL. More than one mismatch was observed in three isolates. For instance, the isolate SRR7750849 was phenotypically resistant to tetracycline and ciprofloxacin yet lacked the *gyrA* point mutation and tetracycline resistance gene.

**Table 4.** Correlation of resistance phenotype and predicted genotype of clinical *C. jejuni* isolates.

| Drug class | No. isolates resistant (%)<br>based on phenotypic<br>testing (n=547) | Resistance gene/point<br>mutation | Cohen's Kappa statistics<br>(Standard Error) |
| --- | --- | --- | --- |
| Quinolones | Ciprofloxacin (168, 30.7%) | *gyrA* T86I (n=163) | 0.98 (0.009) |
|  | Nalidixic acid (167, 30.5%) | *gyrA* T86I (n=163) | 0.98 (0.008) |
| Tetracycline | Tetracycline (286, 52.3%) | *tet*(O) (n=261) | 0.96 (0.01) |
|  |  | *tet*(O/32/O) (n=19) |  |
|  |  | *tet*(O/W/32/O) (n=1) |  |
| Macrolides | Azithromycin (13, 2.4%) | 23S rRNA A2075G (n=10)<br>23S rRNA A2074T (n=1) | 0.91 (0.06) |
|  | Erythromycin (13, 2.4%) | 23S rRNA A2075G (n=10)<br>23S rRNA A2074T (n=1) | 0.91 (0.06) |
| Aminoglycosides | Gentamicin (10, 1.8%) | *aph*(2'')-Ih (n=5)<br>*aph*(2'')-If (n=2) | 0.82 (0.1) |
| Lincosamides | Clindamycin (13, 2.4%) | 23S rRNA A2075G (n=10) | 0.87 (0.07) |

## Discussion

In our study, more than 97% of clinical isolates from Minnesota patients had CDT-producing genes, *cdtA, cdtB*, and *cdtC*. The CDT has been shown to induce DNA damage, which ultimately results in cell death and compromises the integrity of intestinal epithelium (22, 23). However, a study by Abuoun et al. recovered CDT-negative *C. jejuni* from clinical stool samples, albeit infrequently, indicating that additional mechanisms besides CDT can cause clinical symptoms of campylobacteriosis in humans (24). Nonetheless, the prevalence of CDT-producing genes was also high in other studies, suggesting that these genes are widely distributed among *C. jejuni* isolates (25–28).

Only a quarter of the clinical isolates had *flaA* and *flaB*, two flagellin genes responsible for motility in *C. jejuni*. Both functional *flaA* and *flaB* genes are required for the maximum motility in *C. jejuni* (29). Previous studies have shown that non-motile *C. jejuni* was able to colonize chickens, however at a reduced ability or for a short term, indicating that motility is an important virulence factor as it allows *C. jejuni* to migrate and move within the thick mucosal lining of the intestinal tract (30–32). A similar prevalence of *flaA/B* has been reported by Panzenhagen et al., which looked at the distribution of virulence factors in *C. jejuni* sequences deposited in publicly available databases (28). Possible explanations for such a low prevalence provided by Panzenhagen et al. were homologous recombination and intragenic recombination (28). Recombination, which occurs frequently in *C. jejuni*, is thought to be the major source of genetic diversity in *C. jejuni* (33). As a result, *flaA* typing, which is occasionally used to discriminate closely related *C. jejuni* isolates, may not be reliable and accurate (28).

The *virB11* virulence gene is located on the pVir plasmid. In our study, *virB11* and other type IV secretion system genes were found only in 5% of the isolates. According to a study by Wysok et al., the absence of *virB11* in *C. jejuni* led to a slight reduction in HeLa cell invasion, while the absence of the *iam* gene, an invasion-associated marker, significantly reduced cell invasion (34). The prevalence of the *iam* gene was higher than *virB11* in the study by Wysok et al. as well (34). Coupled with the reported low prevalence of *virB11* in *C. jejuni* in chickens and humans (35, 36), the *virB11* gene may not be a critical virulence factor for campylobacteriosis in humans (35, 37).

To perform genome-based prediction of AMR, assembled genomes were screened through a curated ResFinder database. In our study, 29% of clinical isolates carried a *gyrA* point mutation, which confers resistance to quinolones. This level of resistance was similar to the prevalence of quinolone-resistant *C. jejuni* in humans and retail chickens reported by NARMS (6). Five different *gyrA* point mutations were detected. Of these, T86I, which is associated with a high level of fluoroquinolone resistance in *C. jejuni*, was the most frequently observed mutation, consistent with previously reported observations (38). We also detected D90N and P104S mutations, which have previously been detected in clinical isolates (39–41). Interestingly, these mutations were detected together with the T86I mutation.While the D90N mutation is associated with an intermediate level of fluoroquinolone resistance (42), the P104S mutation on its own has been detected in phenotypically sensitive isolates and requires an additional mutation, T86I, to be phenotypically resistant (43). It has been suggested that stepwise accumulation of the *gyrA* point mutation does not lead to an increased level of quinolone resistance (44). Therefore, it is unclear how the acquisition of additional *gyrA* point mutations, P104S and D90N might affect the levels of resistance to quinolones.

We observed a high percentage of isolates carrying beta-lactam resistance genes. Sixteen beta-lactam resistance genes were detected, with more than half of the isolates carrying *bla*_OXA-61_, *bla*_OXA-193_, *bla*_OXA-450_, *bla*_OXA-451_, *bla*_OXA-453_, and *bla*_OXA-489_. *Campylobacter* is known to be intrinsically resistant to beta-lactam antimicrobials, such as penicillin and narrow-spectrum cephalosporins (3). However, the exact mechanisms of beta-lactam resistance in *C. jejuni* are not well defined (3, 45). As such, ampicillin or any other beta-lactam antibiotics have limited application in the treatment of campylobacteriosis (46), and the screening of beta-lactam resistance genes in *C. jejuni* is rarely conducted. However, expanded use of WGS to characterize *C. jejuni* has revealed a widespread presence of beta-lactam resistance genes in *C. jejuni* isolates (47–49). Among the beta-lactam resistance genes detected in the study, *bla*OXA_61_ has been linked to increased resistance to ampicillin in *C. jejuni* (45). Of note, it has been shown that the single nucleotide mutation upstream of *bla*OXA_61_, which modulates the expression of *bla*OXA_61_, might be responsible for ampicillin resistance in *C. jejuni* rather than the presence of the gene itself, as *bla*OXA_61_-carrying *C. jejuni* isolated from poultry was found to be susceptible to ampicillin (50, 51). These findings further support that WGS is a useful tool for understanding resistance modulation mechanisms in addition to identifying novel resistance genes.

More than 50% of the isolates in the study had a tetracycline resistance marker, consistent with observations reported by other studies (47, 52–54). The *tet*(O) was the most frequently detected tetracycline marker; however, two mosaic *tet*(O) genes were detected at a lower frequency. The mosaic *tet* genes are proven to be functional and confer resistance to tetracycline (55). Surveillance of mosaic *tet* genes has been done primarily using PCR-based assays. However, this approach requires knowledge of the target gene sequences, likely to result in an underestimation of *tet* resistance in *C. jejuni*.Implementing WGS to conduct AMR surveillance in foodborne pathogens can improve our understanding of the prevalence and diversity of mosaic tetracycline genes (56).

AST was performed on a subset of the isolates to compare phenotypic and genotypic antimicrobial resistance patterns. Similar to the genotype prediction results based on WGS, 30% of the isolates were resistant to quinolone antibiotics based on phenotypic testing, and a high prevalence of tetracycline resistance (52%) was observed. When resistance phenotype profiles were compared to the genotypic prediction, we observed only 24 discrepancies out of 547 isolates compared. The majority of the observed discrepancies occurred when isolates were phenotypically resistant but lacked the known genetic markers. Notably, three isolates showed multiple discrepancies. For example, one isolate was phenotypically resistant to macrolides, clindamycin, and quinolones, but lacked known point mutations in the 23s rRNA gene and *gyrA* gene. In addition, four isolates were phenotypically resistant to ciprofloxacin, but no point mutation was detected in *gyrA* that could explain the observed phenotype. This result can be attributed to sequencing artifacts or unknown resistance mechanisms in *C. jejuni*. The cmeABC efflux pump plays an important role in antimicrobial resistance in *C. jejuni* by removing various antimicrobial agents (57). Previous studies have shown that an increased expression of *cmeA*, a subunit of the cmeABC operon, led to increased resistance to tetracycline and ciprofloxacin (52, 58). Differential expression of *cmeA* was also observed between ciprofloxacin-susceptible and resistant *C. jejuni* (52), suggesting additional molecular mechanisms conferring resistance to fluoroquinolone antibiotics besides *gyrA* point mutations. Notably, three isolates carried the *tet*(O) gene but were phenotypically susceptible, with their MICs for tetracycline ≤ 0.25 µg/mL. The detection of the *tet*(O) gene in clinical and poultry *C. jejuni* isolates has shown that *tet*(O) can be present on a plasmid rather than located on the chromosome (59, 60). It is plausible that while these three isolates carried the *tet*(O) genes, they were located on a plasmid and were not expressed, resulting in phenotypic susceptibility to tetracycline.

Overall, our results show a high level of concordance between genotypic AMR prediction and phenotypic resistance, consistent with findings from several other studies (52, 61–63). NARMS now employs WGS data to predict AMR in enteric bacteria, including *Salmonella* and *Campylobacter*. Their findings demonstrate that WGS-based AMR prediction correlates well with phenotype data (MICs) for most antimicrobial drugs, supporting the use of WGS as a rapid and reliable method for AMR screening (64). However, standardization of WGS method, use of bioinformatics tools, and AMR reference databases may be warranted to standardize experimental procedures across different laboratories to achieve comprehensive and accurate results.

In conclusion, our study identified a high prevalence of key virulence factors, such as CDT-producing genes, and highlighted the genetic diversity of resistance genes, particularly those associated with beta-lactam resistance. High correlation between genotypic and phenotypic AMR profiles was detected for tetracyclines and quinolones, although discrepancies detected indicate potential gaps in our understanding of resistance mechanisms or limitations with current sequencing technologies. These findings highlight the utility of WGS in enhancing our understanding of AMR and virulence in *C. jejuni*. Overall, incorporating WGS into routine AMR monitoring could improve our ability to detect and manage antimicrobial resistance in foodborne pathogens.

## Acknowledgements

We thank members of the Enterics unit at MDH-PHL for providing laboratory support, including isolation, speciation, and DNA extraction of *C. jejuni* isolates used in this study. We also thank the Sequencing & Bioinformatics Unit at MDH-PHL for performing Illumina sequencing. Lastly, we thank Christy Bennett from Centers for Disease Control and Prevention for reviewing the AST methods.

